# Family History Assessment Significantly Enhances Delivery of Precision Medicine in the Genomics Era

**DOI:** 10.1101/2020.01.29.926139

**Authors:** Yasmin Bylstra, Weng Khong Lim, Sylvia Kam, Koei Wan Tham, R. Ryanne Wu, Jing Xian Teo, Sonia Davila, Jyn Ling Kuan, Sock Hoai Chan, Nicolas Bertin, ChengXi Yang, Steve Rozen, Bin Tean Teh, Khung Keong Yeo, Stuart Alexander Cook, Lori A. Orlando, Saumya Shekhar Jamuar, Geoffrey S. Ginsburg, Patrick Tan

## Abstract

**Background:** Family history has traditionally been an essential part of clinical care to assess health risks. However, declining sequencing costs have precipitated a shift towards genomics-first approaches in population screening programs, with less emphasis on family history assessment. We evaluated the utility of family history for genomic sequencing selection.

**Methods:** We analysed whole genome sequences of 1750 healthy research participants, with and without preselection based on standardised family history collection, screening 95 cancer genes.

**Results:** The frequency of likely pathogenic/ pathogenic (LP/P) variants in 884 participants with no family history available (FH not available group) (2%) versus 866 participants with family history available (FH available group) (3.1%) was not significant (*p*=0.158). However, within the FH available group, amongst 73 participants with an increased family history cancer risk (increased FH risk), 1 in 7 participants carried a LP/P variant inferring a six-fold increase compared with 1 in 47 participants assessed at average family history cancer risk (average FH risk) and a seven-fold increase compared to the FH not available group. The enrichment was further pronounced (up to 18-fold) when assessing the 25 cancer genes in the ACMG 59-gene panel. Furthermore, 63 participants had an increased family history cancer risk in absence of an apparent LP/P variant.

**Conclusion:** Our findings show that systematic family history collection remains critical for health risk assessment, providing important actionable data and augmenting the yield from genomic data. Family history also highlights the potential impact of additional hereditary, environmental and behavioural influences not reflected by genomic sequencing.

## Introduction

Family history is a health assessment tool which can inform risk stratification based on the integration of genetic, environmental and behavioural influences. Traditionally, family history has been used to guide risk assessment of an underlying genetic predisposition in conjunction with a personal history of a medical condition. The identification of a causative genetic variant can inform why disease occurred, highlight potential risks of developing further disease and suggest medical interventions to reduce disease development or progression.^1^ Although family history is a significant indicator for health evaluation, its collection and interpretation can be labour intensive and challenging. Additional difficulties can be encountered when interpreting family history information if collection is incomplete and details are non-specific, or insufficient training is provided to utilise family history information to support clinical decisions.^2^ These challenges are further pronounced when collecting comprehensive family histories for large scale population studies.

With technology advancements and declining costs, genomic sequencing has become increasingly wide-spread, extending into clinical diagnosis and treatment applications. Predicting health risks has expanded from predictive testing within a family where a disease-causing variant is known to be present to analysing a pre-defined set of genes for health risk assessment, driving preventative and personalised medical care at a population level for clinical management. Several screening programs are initiating genomic sequencing for healthy or unselected populations, irrespective of health status or family history.^3–9^ Amongst these population screening programs there is consensus to return genomic results which are medically significant however there is less concordance regarding which genes fall into this category.

The extent to which family history is collected in these initiatives is also variable. Some cohort studies collect a three-four generation pedigree at recruitment,^3,10^ while others gather family history once a clinically significant genomic variant is detected.^6,8,9^ There is emerging evidence that in unselected populations, 48%-60% of individuals identified as having a clinically actionable variant have no associated family history.^8,9,11^ A case control study observed a similar prevalence of Ashkenazi Jewish *BRCA* founder variants amongst individuals with and without breast cancer in their family^12^, and a more recent study found the frequency of clinically actionable variants was similar amongst breast cancer patients who met and did not meet clinical criteria for genetic testing.^13^ These studies have provided evidence to support that testing could be offered to affected and unaffected individuals, with and without an associated increased risk family history.

These results have suggested that in the advent of genome sequencing approaches for large populations as an initial screen, the additional value of family history is debatable.^14^ However, while such studies may suggest that family history is not an accurate tool to detect genomic variants, for many of these studies family history collection time points are not clearly documented or family history is assessed after the detection of a clinically significant variant. Here, we aimed to provide a comprehensive assessment regarding the utility of family history by systematically collecting a three-four generation family history prior to genomic sequencing and comparing the detection of clinically significant genomic variants in cancer predisposition genes amongst 1750 healthy participants with no known pre-existing medical conditions.

## Methods

### Study design and participants

This cohort study conducted in Singapore examined the frequency of clinically significant variants in participants that reported to have an increased family history of cancer and participants unselected for risk or at average risk according to their family history. The participants were recruited for a prospective institutional review board-approved Biobank or SingHeart study (https://clinicaltrials.gov/ct2/show/study/NCT02791152) conducted at the National Heart Centre Singapore between August 2014 to December 2018. Details of participant recruitment and methods of both Biobank and SingHeart have been previously described.^15^ Briefly, volunteers with no known pre-existing health conditions over 16 years of age were recruited in response to a research advertisement in the local paper in 2014. They consented to a detailed medical screen and a genetic screen using whole genome sequencing technology. MeTree (an online family history collection tool) was incorporated into the recruitment process in 2016 to systematically collect family history. Prior to this family history was not collected at recruitment. All participants included in this study were asymptomatic as ascertained after their health screen at recruitment and none reported previous diagnosis of cancer.

### Family history collection

For participants where family history was collected, participants were notified prior to their initial recruitment appointment to gather medical information from their family members. At their recruitment appointment, family history was collected using an online family history tool (MeTree)^16^ which prompts for a range of conditions such as cancer and heart conditions and has current U.S. clinical guidelines incorporated to create personalised risk reports for patients and their providers.

### Risk assessment based on family history

Each family history documenting a presence of cancer was assessed by the clinical genetics team in accordance with clinical testing criteria guidelines, National Comprehensive Cancer Network (NCCN) Genetic/ Familial High-Risk Assessment Breast and Ovarian (Version 3.2019) ^17^ and Genetic/Familial High-Risk Assessment:Colorectal (Version1.2018)^17,18^ or supplemented by an organ-specific international guideline to determine the risk of developing cancer.^19,20^ In cases where the familial risk was unclear because of incomplete information pertaining to cancer type, age of diagnosis or disease progression, the family pedigree was reviewed in further detail by the clinical genetics team taking into consideration participant age and number of family members until a consensus of risk was reached.

### Cancer genes for analysis

A gene list associated with cancer development was devised from genes well-studied in the literature and ranged from some to strong evidence for cancer susceptibility. This gene list was subsequently compared with several databases such as OMIM (Online Mendelian Inheritance in Man)^21^ and ClinGen (Clinical Genome Resource).^22^ Twenty-five of these comprise the American College of Medical Genomics and Genetics (ACMG) 59 secondary finding gene list. (Supplementary Table 1).

### Genomic sequencing and classification

DNA was extracted from a donated blood sample and WGS was performed with a third party provider using the Illumina HiSeq X platform under standard protocols. Data was returned in the form of FASTQ files and analysed using an in-house bioinformatics pipeline as previously described.^15^

Variants occurring in the customised cancer gene panel were filtered by frequency against a population-matched database and selected according to their classification for both exonic and intronic regions in ClinVar^23^ or disruption to protein function and curated according to ACMG guidelines^24^.

Haploinsufficiency for each gene was assessed by literature review and recommendation in ClinGen (accessed until May 2019)^22^. Consensus for the variant classification was obtained by discussion amongst genetics specialists. For each variant classified as either likely pathogenic or pathogenic, the QC metrics and corresponding BAM files were then visually inspected for confirmation. For variants where the QC metrics and/or presence in BAM was ambiguous, these were then validated by Sanger sequencing.

### Statistical analysis

Relative risk (RR) was calculated as specified by Altman *et al.* 1991.^25^ All statistical tests were two-tailed and a p-value less than 0.05 was considered statistically significant.

## Results

### Cohort descriptions

We performed targeted genome analysis of 1750 participants. Of these 884 did not have family history available (FH not available group) as they were recruited prior to the integration of family history and 866 had family histories recorded (FH available group). Within the FH available group, testing guidelines criteria revealed 73 families (8.4%) to be at increased risk of developing cancer (increased FH risk). There were 793 participants found not to be at increased risk based on their family history (average FH risk) (Figure 1).

**Figure 1:**
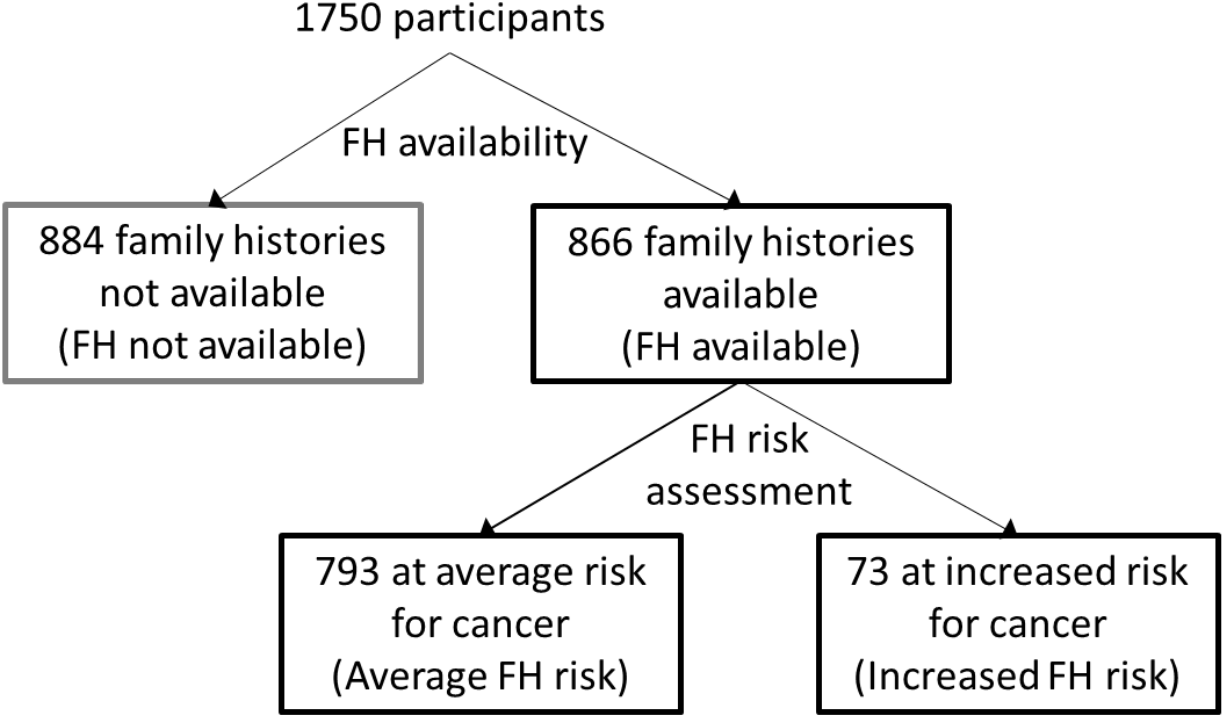
Family history assessment overview. The participant cohort was divided according to the availability of family history – FH not available and FH available. The FH available group was further divided according to assessment of family history cancer risk – Average FH risk and Increased FH risk.

Apart from gender in the increased FH risk group, the characteristics between all groups were similar. The mean age, gender and ethnicity for each group is represented in Table 1. An overview of the cancers reported in each family is provided in Supplementary Figure 1.

**Table 1.**
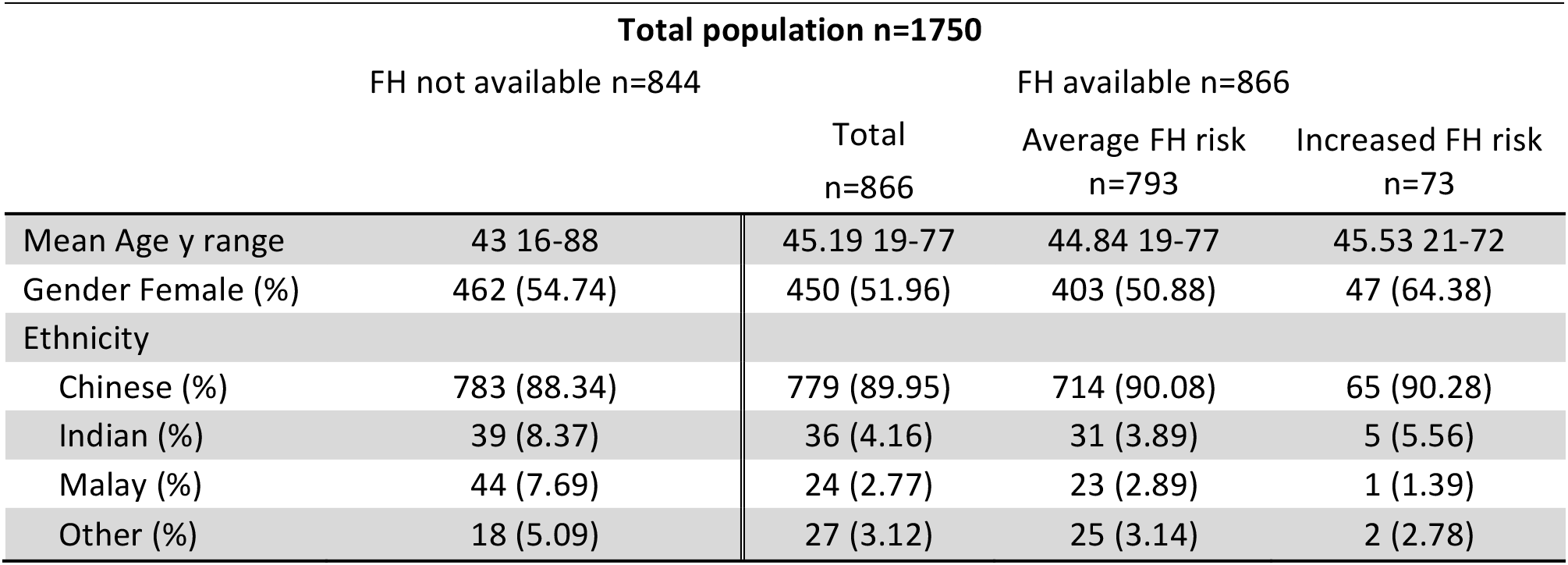
Characteristics of cohorts.

### Total population analysis

Overall, there were 45 likely pathogenic/ pathogenic (LP/P) variants detected amongst the 1750 participants and these were grouped according to family history availability and risk assessment (Figure 2). Amongst the FH not available group, 2% (17/884) or 1 in 52 participants were found to carry a LP/P variant. (Table 2).Seventeen LP/P variants were found in 14 cancer genes, four of these occurred in three genes from the ACMG secondary finding list (2 in *BRCA2*, 1 in *MSH6* and 1 in *TP53*) and two participants carried the same variant (Supplementary Table 2).

**Table 2.**
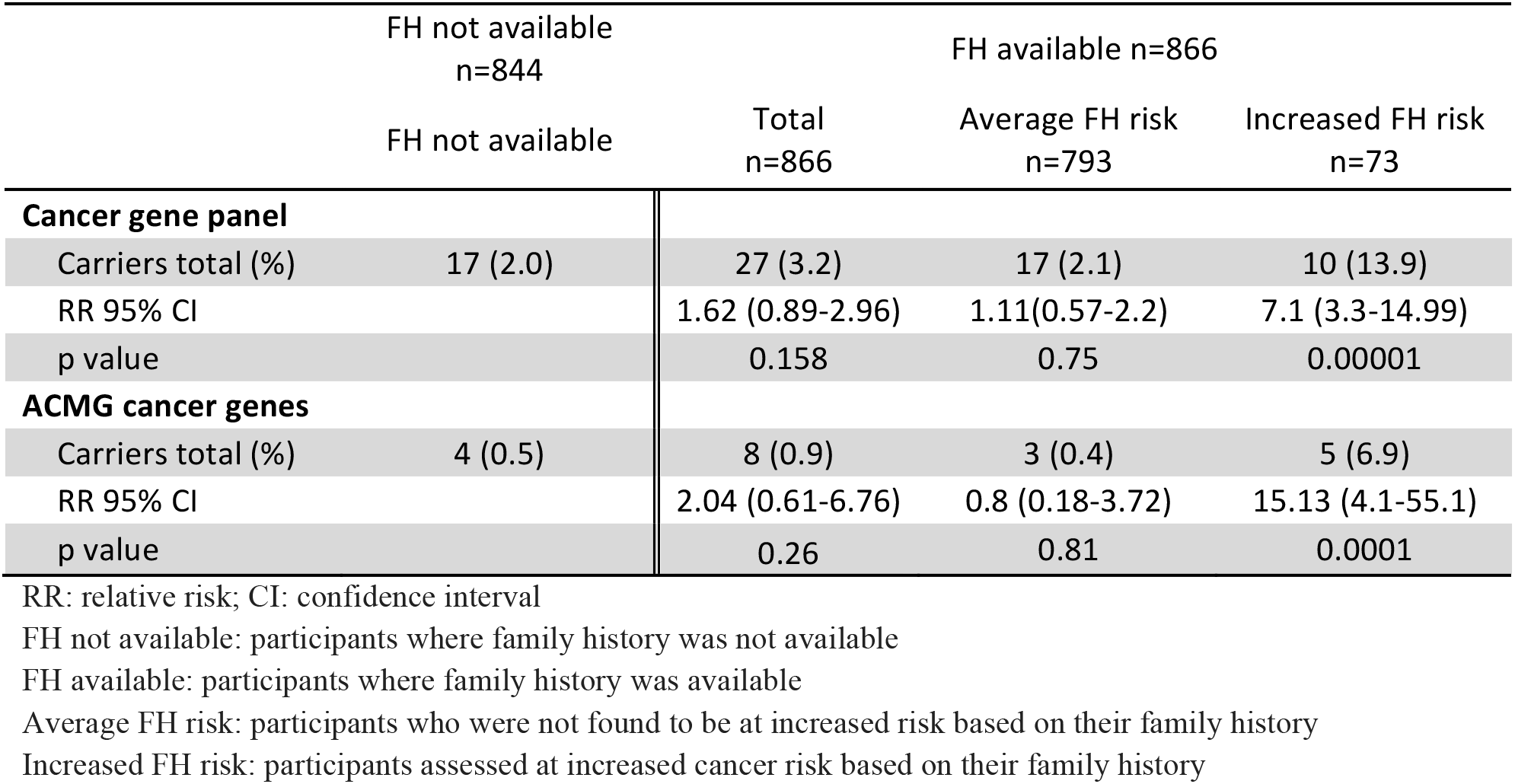
Comparison of LP/P variants identified in FH not available and available groups.

**Figure 2.**
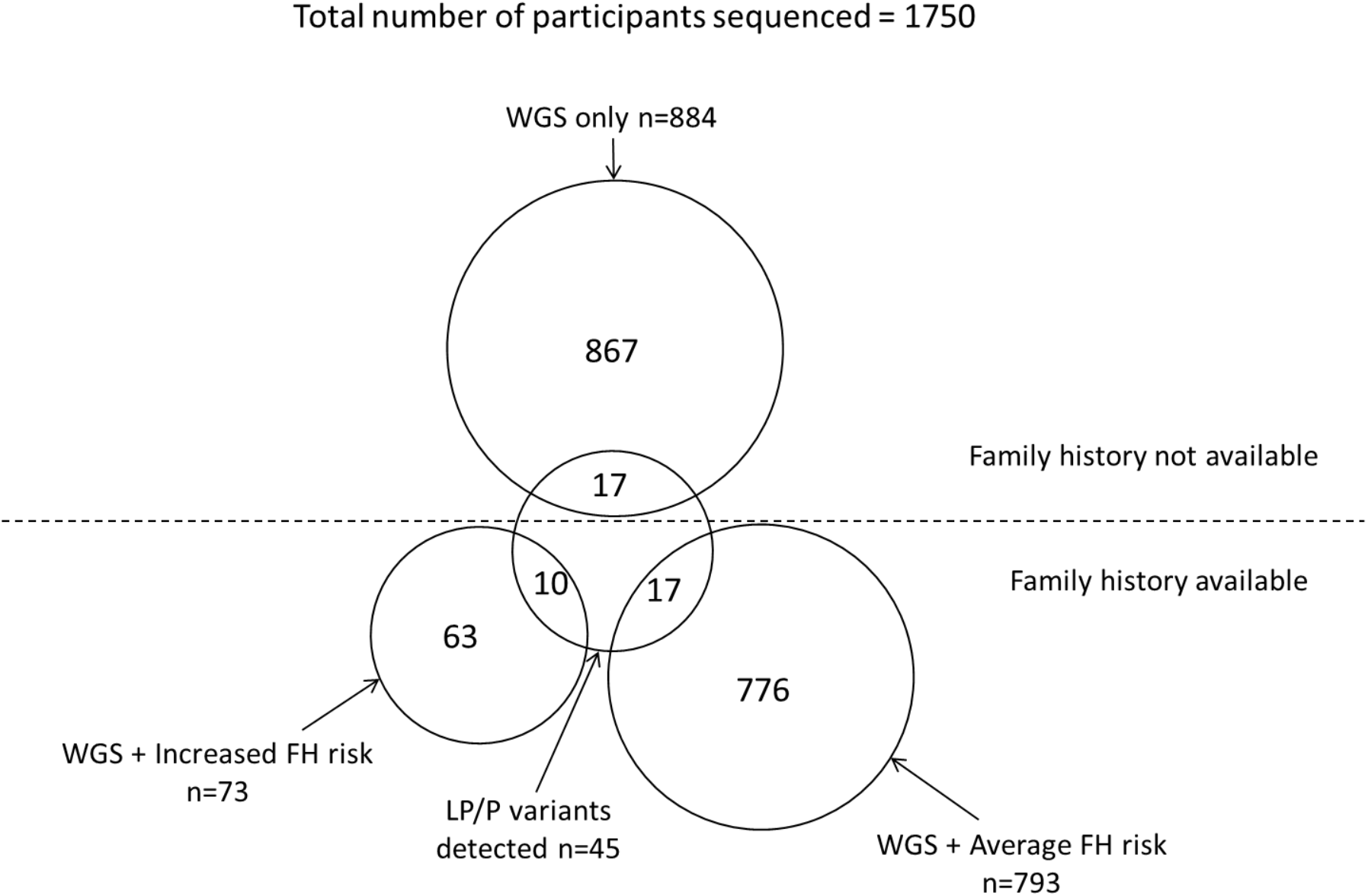
Number of LP/P variants detected. The inner circle represents the number of individuals with LP/P variants detected from each of the three cohorts – the FH not available cohort where only WGS was available, the WGS + increased FH risk for cancer cohort and the WGS + FH average risk for cancer cohort. The outer circles represent the number of individuals who were not found to carry a LP/P variant.

In the FH available group, 3.1% (27/866) or 1 in 32 participants were found to have a LP/P variant. Twenty-seven LP/P variants were found in 15 cancer genes (Table 2) and eight of these occurred in three genes from the ACMG secondary finding list (2 in *BRCA1*, 5 in *BRCA2* and 1 in *MSH2*) (Supplementary Table 2).

The frequency of clinically actionable variants detected between the FH not available and available groups was not statistically significant (*p*=0.158). No participants were found to be homozygous or compound heterozygous for an autosomal recessive condition, or were carriers of one of the seven autosomal recessive genes and had an associated family history from either the FH not available or available groups.

### FH available cohort analysis

Once ascertained for cancer risk according to family history in the FH available group, the variants with clinical significance were more frequent in the increased FH risk group compared to the FH not available group (Table 2) and average FH risk group (Table 3). Amongst the increased FH risk group, 13.9% (10/73) or 1 in 7 unrelated participants were found to have a LP/P variant. Ten LP/P variants were identified occurring in five cancer genes. Of these 10 variants, five were found in three of the genes in the ACMG secondary findings list (2 in *BRCA1*, 2 in *BRCA2* and 1 in *MSH2*). The other five were in *ATM (2)*, *AXIN2 (1), RAD50 (1) and SUFU (1)* (Supplementary Table 2). Furthermore, the probability of carrying a LP/P variant and having an increased risk family history (10/27) was three times higher (relative risk (RR) 3.87, 95% confidence interval (CI) 2.17-6.91, *p*<0.00001) than having an increased risk family where no LP/P variant was found (63/839) (Table 4).

**Table 3.** Comparison of LP/P variants identified within the FH available group.

|  | FH available n=866 |  |
| --- | --- | --- |
|  | Average FH risk<br>n=793 | Increased FH risk<br>n=73 |
| **Cancer gene panel** |  |  |
| Carriers total (%) | 17 (2.1) | 10 (13.9) |
| RR 95% CI |  | 6.39(3.04-13.4) |
| Fisher's exact test p value |  | 0.00001 |
| **ACMG cancer genes** |  |  |
| Carriers total (%) | 3 (0.4) | 5 (6.9) |
| RR 95% CI |  | 18.1(4.41-74.2) |
| Fisher's exact test p value |  | 0.0002 |

RR: relative risk; CI: confidence interval
Average FH risk: participants who were not found to be at increased risk based on their family history
Increased FH risk: participants assessed at increased cancer risk based on their family history

**Table 4.** LP/P variants detected according to family history analysis.

|  | LP/ P variants<br>detected n=45 | No variant detected<br>n=1705 |
| --- | --- | --- |
| FH not available group | 17 (40%) | 867 (51%) |
| FH available group | 27 (60%) | 839 (49%) |
| Average FH risk | 17 (63%) | 776 (92.5%) |
| Increased FH risk | 10 (37%) | 63 (7.5%) |

Of the 10 causative variants detected in the increased FH risk group, eight participants carried a clinically actionable variant where the association with their family history was well established. For example, participant 35 reported a family history suggestive of Lynch syndrome^26^ and carried a *MSH2* variant. However, there was one participant with an *AXIN2* variant and a family history of breast cancer and evidence regarding this association is only emerging (Supplementary Table 3).

Amongst the 793 participants in the average FH risk group, 2.1% (17/793) or 1 in 47 unrelated participants were found to carry a LP/P variant. There was no significant difference between the average FH risk and FH not available groups and detection of clinically actionable variants (*p*=0.75) (Table 2). Seventeen LP/P variants were found in 11 cancer genes. Of these, eight carriers reported a family history of cancer, however, as the age of diagnosis was older or unknown they did not meet the pre-specified clinical testing criteria for increased risk (Supplementary Table 3). Three of these variants occurred in *BRCA2* from the ACMG secondary findings gene list (Supplementary Table 2).

## Discussion

As family history has been long understood to play a vital role in targeting underlying genetic causes, we conducted an in-depth assessment of family history and genomic data in a population genomic screening study setting. Amongst a cohort of 1750 participants who had undergone genome sequencing, 866 family histories of three- four generations were collected. We did not find any significant difference in the number of cancer gene carriers between the two groups with and without available family history. Where family history was available, 73 participants were at increased risk of developing cancer and 1 in 7 participants carried an autosomal dominant LP/P variant which was a six-fold increase when compared to the FH average risk group (1 in 47) and a seven-fold increase when compared to the FH not available group. This threshold was further pronounced when selecting for the 25 cancer genes in the 59 ACMG gene panel amongst the increased FH risk (1 in 14 or 6.9%) versus the average FH (1 in 264 or 0.4%) and FH not available (1 in 221 or 0.5%) groups. The prevalence of ACMG cancer gene LP/P variants in our increased FH risk group was also considerably higher than other reported studies that assessed the presence of pathogenic variants in the 59 ACMG gene list of unselected populations ranging from 1.5%^27^ to 2.7%^7^ and 1.6% in an ethnically similar cohort.^28^

Large scale population screening programs have been initiated globally to optimise the utility of genomic data. The analysis of genomic data has unquestionably led to a greater understanding of the genetic basis of health and disease. However, the collection of additional clinical and personal data and the return of genomic results continues to be refined in light of cost effectiveness, clinical utility and the psychological impact of receiving such results. Some recent cohort studies have suggested that family history is not a useful tool for identifying carriers of monogenic conditions, as at least half of the carriers detected in their unselected populations did not present with a corresponding increased risk family history, nor would have met eligibility criteria for genetic testing.^8,9,11^ We also detected carriers of cancer syndromes that did not meet testing guidelines according to their family history (17 participants), half of which had no family history of cancer. Possible explanations include reduced penetrance or a milder phenotype of the disease-causing variant in absence of other environmental or genetic risk factors not present in these families or that the identified variant is *de novo*. It is also possible that the family histories are incomplete and that further relevant family information could be revealed.^8^

In contrast, by integrating family history assessment, our findings indicated that selecting participants according to their family history for genomic testing significantly increased the detection of carriers for cancer syndromes. Therefore, the traditional triaging of participants by family risk assessment in our study appears to be a cost-effective alternative in contrast to genome screening unselected populations to increase the detection of clinically actionable variants. This would be particularly relevant in a resource-constricted environment where genomic testing may not be available to the wider population. Furthermore, family history provides a useful tool to frame expectations when counselling about the likelihood of detecting disease-causing variants. There is also evidence supporting that asymptomatic individuals are more likely to adhere to recommended screening programs with experiential knowledge of the condition they are at risk of developing, which frequently derives from family history over time.^29,30^

There was also a significant proportion of participants that reported an increased risk of cancer where no clinically significant genomic variants were detected. Possible genetic explanations include copy number variants, the involvement of genes outside our customised gene panel or a combination of genetic factors contributing to oligogenic inheritance. We also adopted strict classification criteria to annotate the variant pathogenicity and it is likely that there are many variants of unknown significance present in these participants which over time could be identified as disease-causing. Health is understood to be influenced by multiple factors including social circumstances, environmental exposures, behavioural patterns and healthcare systems, with genetic predisposition only contributing 30%.^31^ Expanding beyond the focus of monogenic disease risk, family history has the potential to capture interactions between hereditary, environmental and behavioural factors that are present within families.^32,33^ As such, these participants would still meet cancer surveillance recommendations based on their family history which would not have been evident if genomic sequencing was initiated as a health screen without evaluation of family history as well.

There was one participant in the increased FH risk group was found to carry a LP variant in *AXIN2* and a family history of breast cancer*. AXIN2* has more recently been described to be associated with colorectal cancer^34^ and therefore the association with breast cancer is not well understood. However, even with the removal of *ATM* and *AXIN2* from the increased FH risk group, there is still a five-fold increase (RR 5.8, 95% CI, 2.6-12.4, *p* < 0.0001) in detecting LP/P variants in participants with an increased risk family history. Over time we may learn that theis variant is unrelated to the family history or instead, that it corresponds to the expansion of currently understood genotype-phenotype correlations.

Our study was limited by the relatively small number of families found to have a significant family history risk of cancer in comparison to the average risk group and the FH not available group. The assessment of family history relies on the accuracy of the information provided. Even though there are studies demonstrating family history recollection is reliable^35,36^, as this information is self-reported it could be incomplete or imprecise, thus impacting the reliability of risk assessment. The assessment of condition-specific risk using established guidelines can be time-consuming and challenging if family history information is incomplete. Further work could involve modifying the risk assessment criteria to optimise how much family information is required as triage for the assessment of pathogenic variants.

This study has illustrated the utility of documenting and assessing detailed family history to increase the detection of LP/P variants using genomic sequencing in a healthy cohort. In a clinical setting, these findings provide a practical tool to frame the likelihood of detecting a clinically significant variant, manage expectations and assist with decision-making when genomic sequencing is offered. At a population screening level we propose that family history is an effective screening tool to triage individuals that would most benefit and an integral step towards extending precision medicine applications to precision testing. Furthermore, our findings indicate that family history can assess for personal disease risk beyond genetic factors as evidenced by participants with a family history yet no concerning genetic variant found. In conclusion, we have demonstrated that comprehensive family history collection continues to have a significant role in this genomic era.

## Acknowledgments

We thank the volunteers who participated in both Biobank and SingHEART research studies, Siew Ching Kong and Pei Yi Ho, National Heart Centre Singapore, for their roles in recruitment and participant contact and the assistance of Nelly Chai Bin Siew, Duke-NUS Singapore and Hui Hoon Chua, Biomedical Research Council, Singapore with MeTree.

