## Supplementary Information for "Family History Assessment Significantly Enhances Delivery of Precision Medicine in the Genomics Era"

### Supplementary Material

Table 1. Gene list
Figure 1. Cancer prevalence amongst the 73 increased FH risk participants
Table 2. LP/P variants found in the FH not available and FH available groups
Table 3. LP/P variants and associated family history in the increased and average risk cohorts

**Supplementary Table 1. Cancer gene panel**

The gene list comprises 95 genes associated with 28 hereditary cancer syndromes and 18 organ specific cancer types. Twenty-five of these are part of the 59 medically actionable dominant disease genes for which American College of Medical Genomics and Genetics (ACMG)[24] recommends disclosure. Seven genes are associated with autosomal recessive inheritance.

| **Gene, MIM#** | | | | |
|---|---|---|---|---|
| *AKT1*, 164730 | *CEBPA*, 116897 | *LZTR1*, 600574 | *PDGFRA*, 173490 | ***SDHB*, 185470** |
| *ALK*, 105590 | *CEP57*, 607951 | *MAX*, 154950 | *PHOX2B*, 603851 | ***SDHC*, 602413** |
| ***APC*, 611731** | *CHEK2*, 604373 | ***MEN1*, 131100** | *PIK3CA*, 171834 | ***SDHD*, 602690** |
| *ATM*, 607585 | *DICER1*, 606241 | *MET*, 164860 | ***PMS2*, 600259** | *SLX4*, 613278 |
| *AXIN2*, 604025 | *DIS3L2*, 614184 | *MITF*, 156845 | *POLD1*, 174761 | ***SMAD4*, 600993** |
| *BAP1*, 603089 | *EGFR*, 131550 | ***MLH1*, 120436** | *POLE*, 174762 | *SMARCA4*, 603254 |
| *BARD1*, 601593 | *EPCAM*, 185535 | *MLH3*, 604395 | *POT1*, 606478 | *SMARCB1*, 601607 |
| *BLM*, 210900 | *EZH2*, 601573 | *MRE11A*, 600814 | *PRKAR1A*, 188830 | *SMARCE1*, 603111 |
| ***BMPR1A*, 601299** | *FANCA*, 607139 | ***MSH2*, 609309** | *PTCH1, 601309* | ***STK11*, 602216** |
| ***BRCA1*, 113705** | *FANCC*, 613899 | *MSH3*, 600887 | ***PTEN*, 601728** | *SUFU*, 607035 |
| ***BRCA2*, 600185** | *FH*, 136850 | ***MSH6*, 600678** | *RAD50*, 604040 | *TERT*, 187270 |
| *BRIP1*, 605882 | *FLCN*, 607273 | ***MUTYH*, 604933** | *RAD51C*, 602774 | *TMEM127*, 613403 |
| *BUB1B*, 602860 | *GALNT12*, 610290 | *NBN*, 602667 | *RAD51D*, 602954 | ***TP53*, 191170** |
| *CDC73*, 607393 | *GATA2*, 137295 | *NF1*, 162200 | ***RB1*, 614041** | ***TSC1*, 605284** |
| *CDH1*, 192090 | *GPC3*, 300037 | ***NF2*, 607379** | ***RET*, 164761** | ***TSC2*, 191092** |
| *CDK4*, 123829 | *HOXB13*, 604607 | *NKX2-1*, 600635 | *RHBDF2*, 614404 | *TSHR*, 603372 |
| *CDKN1B*, 600778 | *HRAS*, 190020 | *NTHL1*, 602656 | *RUNX1*, 151385 | *TSC*, 608537 |
| *CDKN1C*, 600856 | *KIF1B*, 605995 | *PALB2*, 610355 | *SDHA*, 600857 | ***VHL, 60837*** |
| *CDKN2A*, 600160 | *KIT*, 164920 | *PALLD*, 608092 | ***SDHAF2*, 613019** | ***WT1*, 607102** |

Genes bolded are in the 59 ACMG reportable gene list
Genes underlined are associated with autosomal recessive inheritance

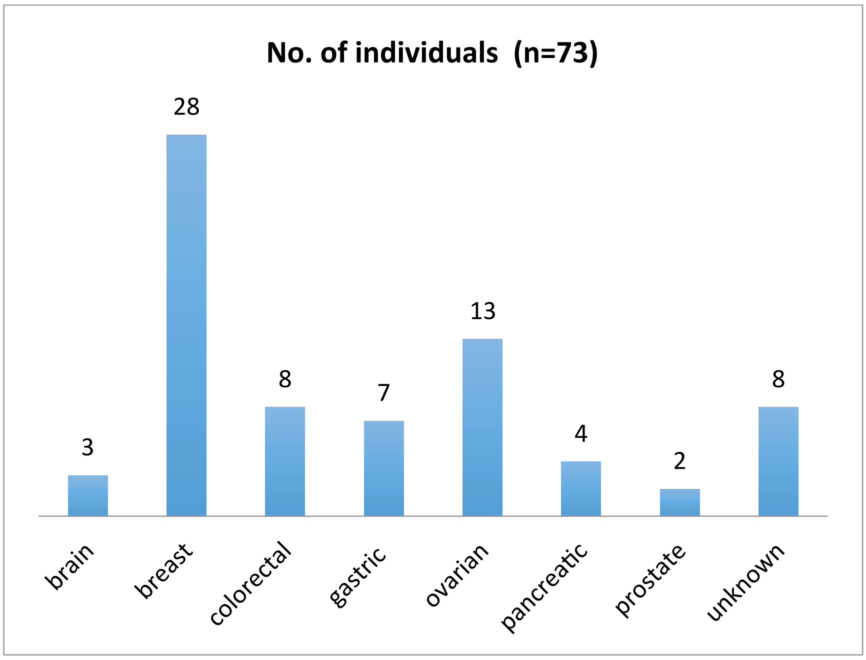


**Supplementary Figure 1. Cancer prevalence amongst the 73 increased FH risk participants**

To provide an overview of cancer risk in the increased FH cohort, these families were grouped by the cancer most pre-dominant in the family, either due to its frequency or an earlier-than-expected age-of-diagnosis. The most commonly reported cancer in the family was breast cancer. Some participants indicated multiple family members with early-onset cancer e.g. aged 30s and 40s but were unaware of the cancer type, forming the unknown category.

**Supplementary Table 2 – LP/P variants found in the FH not available and FH available groups**

| | **Total population n=1750** | | | |
|---|---|---|---|---|
| | FH not available n=884 | FH available n=866 | | |
| **Gene** | | Total n=866 | Average FH risk n=793 | Increased FH risk n=73 |
| *ATM* | 1 | 4 | 2 | 2 |
| *AXIN2* | - | 1 | - | 1 |
| *BLM* | - | 1 | 1 | - |
| ***BRCA1*** | - | 2 | - | **2** |
| ***BRCA2*** | **2** | **5** | **3** | **2** |
| *BRIP1* | 2 | 1 | 1 | - |
| *CDKN1B* | 1 | - | - | - |
| *CDKN2A* | 1 | - | - | - |
| *DICER1* | - | 1 | 1 | - |
| *GPC3* | - | 1 | 1 | - |
| *LZTR1* | - | 2 | 2 | - |
| *MAX* | 1 | - | - | - |
| ***MSH2*** | - | **1** | - | **1** |
| ***MSH6*** | **1** | - | - | - |
| *NF1* | 1 | - | - | - |
| *RAD50* | 2 | 2 | 1 | 1 |

| | | | | |
|---|---|---|---|---|
| *RAD51C* | 2 | 1 | 1 | - |
| *RAD51D* | 1 | 3 | 3 | - |
| *SDHA* | 1 | - | - | - |
| *SUFU* | - | 1 | - | 1 |
| ***TP53*** | **1** | - | - | - |
| *XRCC2* | - | 1 | 1 | - |
| **Cancer gene panel** | | | | |
| Carriers total % | 17 (2.0 | 27 (3.2) | 17 (2.1) | 10 (13.9) |
| **ACMG cancer genes** | | | | |
| Carriers total % | 4 (0.5 | 8 (0.9) | 3 (0.4) | 5 (6.9) |

Genes bolded are in the 59 ACMG reportable gene list

#### Supplementary Table 3. LP/P variants and associated family history in the increased and average risk cohorts

| Family ID | Gene variant | Family history of cancer |
|---|---|---|
| **Increased FH risk** | | |
| **8** | *ATM* c.8224_8225delAA p.Asn2742HisfsTer4 | Paternal uncle dx colorectal cancer, another paternal uncle dx lung cancer, paternal grandmother dx colorectal cancer |
| **9** | *AXIN2* c.1614_1642del p.R538fs | Father dx heart attack 47y, bypass 48y, heart attack 50y and d. heart failure 65y, Mother dx Parkinson 64y, maternal grandmother dx esophageal cancer d.70y, maternal grandfather dx lymphatic cancer d.71y, maternal aunt 1 dx breast cancer 45y, maternal aunt 2 dx nipple cancer 40y, paternal aunt 2 dx breast cancer d.45y, paternal uncle dx mild stroke 50y |
| **2** | *ATM* c.A7702 p.R2568X | Sister dx breast cancer 50s, maternal uncle dx stomach cancer 40s d.40s, paternal aunt dx unknown cancer d.75y, paternal grandfather dx unknown cancer d.50s |
| **11** | *BRCA1* c.C4231T p.Q1411X | Mother dx ovarian cancer 60s, father dx prostate cancer 50y |
| **37** | *BRCA1* c.2585dupA p.N862fs | Sister dx stomach cancer d.37y, another sister dx ovarian cancer, maternal aunt dx unknown cancer d.65y |
| **54** | *BRCA2* c.4337delT p.I1446fs | Maternal aunt dx ovarian cancer 60s and adenomas, maternal grandmother dx adnenomas |
| **79** | *BRCA2* c.7377_7380del p.K2459fs | Mother dx unknown cancer 40s, sister dx breast cancer 40s |
| **35** | *MSH2* c.G1102T p.E368X | Father dx colorectal cancer 30s, pancreatic cancer 60s, sister dx ovarian cancer 20s, uterine cancer 30s, paternal grandmother dx liver cancer 40s |

| | | |
|---|---|---|
| **58** | *RAD50* c.2157dupA p.L719fs | Father dx liver cancer 40s d.40s, mother dx thyroid disorder, maternal grandmother dx lung cancer 20s d.20s, maternal grandfather dx unknown cancer 40s d.40s, paternal grandmother dx breast cancer 50s d.50s |
| **6** | *SUFU* c.65dupC p.A22fs | Daughter dx brain cancer in infancy |
| **Average FH risk** | | |
| **176** | *ATM* c.8545C>T p.R2849X | Father d. heart failure 80y, sister dx breast cancer, brother dx hypertension |
| **67** | *ATM* c.8434_8435del p.S2812fs | Father dx hypertension 60s, high cholesterol and heart attack d.70s, lung disease 80s, mother dx hypertension 50s and high cholesterol 60s, maternal grandmother dx colorectal cancer and hypertension 50s, maternal grandmother d. lung disease 80s, paternal grandmother dx lung disease d.60s |
| **53** | *BLM* c.1290_1291InsATCAGGCCTCCATAG | Mother dx slight stroke 70s d.85y, daughter dx thyroid issues 25y |
| **15** | *BRCA2* c.9684delT p.S3228fs | Father dx high blood sugar 58y, paternal grandfather dx. lung cancer 50s d.50s, paternal aunt dx diabetes 50s, P uncle dx colorectal cancer/liver cancer 60s |
| **767** | *BRCA2* c.5574_5577del p.T1858fs | Father dx heart disease adolescence and hypertension 60s d.60s, mother dx hypertension, diabetes and diabetic kidney disease 70s, sister dx hypertension 60s, maternal grandmother dx breast cancer 60s |
| **750** | *BRCA2* c.7805+3A>C | Father dx prostate issues and heart attack 60s, mother dx thyroid disease 50s and hypertension 60s, maternal grandmother d. stroke 60s, PGM d. lung disease 80s, paternal aunt dx Parkinson disease 70s |
| **637** | BRIP1 c.1343G>A p.W448X | Father dx hypertension, mother dx diabetes d.55y, sister 2 and 3 dx diabetes |
| **169** | *DICER1* c.1353_1360delp.R451fs | Father dx hypertension 70s and lung cancer 80s, mother dx hypertension, obesity and rhematoid arthritis 70s, maternal aunt 3 dx colorectal 80s d.80s, P aunt 1 d. stroke 60s, P aunt 3 dx breast cancer 60s d.60s |
| **2** | *GPC3* c.67C>T p.Q23X | Father dx hypertension 40s, paternal grandmother d. heart attack 60s |
| **76** | *LZTR1* c.1018C>T p.R340X | Mother dx high cholesterol 70s, Maternal aunt dx ischaemic stroke 70s |
| **81** | *LZTR1* c.465C>G p.Y155X | Father dx hypertension 40s and heart disease 60s, mother dx high cholesterol and diabetes 50, maternal grandmother dx diabetes 80s, paternal grandmother |

| | | |
|---|---|---|
| | | dx liver cancer 70s d.80s, paternal uncle dx hypertension 60s |
| **454** | *RAD50* c.2157delA p.L719fs | Father dx COPD, mother dx cervical cancer 50s, maternal uncle dx prostate cancer 50s, paternal grandmother dx diabetes d. diabetes-related complications 80s, paternal grandfather dx unknown cancer d.90s, paternal uncle dx diabetes |
| **281** | *RAD51C* c.390dupA p.G130fs | Maternal grandfather d. unknown cancer 50s, brother dx unknown liver condition 30s |
| **662** | *RAD51D* c.331_332insTA p.K111fs | Father dx with hypertension, mother dx with diabetes |
| **118** | *RAD51D* c.331_332insTA p.K111fs | Father dx heart attack 79y, mother dx hypertension and diabetes, sister 1 dx hypertension, sister 2 dx unknown cancer 63y |
| **671** | *RAD51D* c.331_332insTA p.K111fs | Father dx bladder cancer 65y and stroke 80y d.85y, mother dx transient ischemic attack 93y, brother dx hypertension and colorectal cancer 76y d.76y, sister dx fibroids and had thyroidectomy |
| **64** | *XRCC2* c.280dupA p.T94fs | Father dx ?lung/throat cancer 50s d.59y |

dx:diagnosis, d.: died, y:year
